# Ultradian rhythms of CRH^PVN^ neuron activity, behaviour and stress hormone secretion

**DOI:** 10.1101/2024.06.30.601439

**Authors:** Shaojie Zheng, Caroline M. B. Focke, Calvin K. Young, Isaac Tripp, Dharshini Ganeshan, Emmet M. Power, Daryl O. Schwenke, Allan E. Herbison, Joon S. Kim, Karl J. Iremonger

**Affiliations:** Centre for Neuroendocrinology and Department of Physiology, University of Otago, Dunedin, New Zealand.; Department of Physiology, Development and Neuroscience, University of Cambridge, United Kingdom.

**Author notes:** denotes equal contribution (Shaojie Zheng, Caroline M. B. Focke). Corresponding author: Karl J. Iremonger, **Email:**. **Author Contributions:** Designed research: SZ, CMBF, KJI. Performed research: SZ, CMBF, IT, DG, EMP, JSK. Contributed reagents/analytic tools: CKY, DOS, AEH, JSK. Analyzed data: SZ, CMBF, CKY, IT, DG, EMP, JSK, KJI. Supervised project: DOS, AEH, KJI. Wrote the paper: SZ, CMBF, KJI. **Competing Interest Statement:** Authors disclose no competing interests. **Classification:** Biological Sciences; Neuroscience.

**Keywords:** CRH, stress, ultradian, behaviour, corticosterone

## Abstract

The stress axis is always active, even in the absence of any threat. This manifests as hourly pulses of corticosteroid stress hormone secretion over the day. Corticotropin-releasing hormone neurons in the paraventricular nucleus of the hypothalamus (CRH^PVN^) control both the neuroendocrine stress axis as well as stress-associated behaviours. However, it is currently unclear how the resting activity of these neurons is coordinated with both spontaneous behaviour and ultradian pulses of corticosteroid secretion. To investigate this, we performed fiber photometry recordings of CRH^PVN^ neuron activity in *Crh-Ires-Cre* mice and a newly generated line of *Crh-Ires-Cre* rats. In both mice and rats, CRH^PVN^ neurons displayed an ultradian rhythm of activity with reoccurring upstates of activity approximately once per hour over the 24-hour day. Upstates in activity were coordinated with increases in animal activity/arousal. Chemogenetic activation of CRH^PVN^ neurons was also sufficient to induce behavioural arousal. In rats, increases in CRH neural activity preceded some pulses of corticosteroid secretion but not others. Thus, while CRH^PVN^ neurons display an ultradian rhythm of activity over the 24-hour day that is coordinated with behavioural arousal, the relationship between CRH^PVN^ activity and pulses of corticosteroid secretion is not one-to-one.

**Significance Statement:** Ultradian rhythms are present in many biological systems, however, the underlying neural mechanisms responsible for generating these rhythms is unclear. Corticotropin-releasing hormone (CRH) neurons control corticosteroid levels in the body as well as arousal and behaviour. Here we show that CRH neurons exhibit a pronounced ultradian rhythm in neural activity over the 24-hour day. CRH neural activity was highly correlated with, and predictive of, behavioral arousal. However, CRH neural activity was not as well correlated with pulses of corticosteroid secretion. Together, these data revel the patterns of CRH neuron activity over the 24-hour day and their relationship with behaviour and stress hormone secretion.

## Introduction

Ultradian rhythms are common in biological systems and are observed in unicellular organisms through to vertebrates. In mammals, ultradian rhythms exist in many systems and result in oscillating gene expression, hormone secretion and behavior over the 24-hour day^1,2^. Indeed, pronounced ultradian rhythms are observed in arousal, locomotion, body temperature and stress hormone secretion^1–7^. Current evidence suggests that ultradian oscillations in different systems are generated by distinct neural circuits which employ unique neural mechanisms to generate these rhythms.

The hypothalamic-pituitary-adrenal (HPA) axis shows a prominent ultradian rhythm of activity^8–11^. This is characterized by pulses of corticosteroid secretion that occur approximately once every 60-90 minutes^7^. Ultradian rhythms in the HPA axis are critical for normal physiological function in mammals and can entrain the expression of glucocorticoid-responsive genes throughout the body and brain^12^. Disrupting these rhythms can disrupt numerous aspects of physiology ranging from stress axis responsiveness to cognitive function and metabolism^11,13^. Despite the importance of ultradian HPA rhythms, the mechanisms which generate them are still contentious.

Corticotropin-releasing hormone neurons in the paraventricular nucleus of the hypothalamus (CRH^PVN^) control the HPA axis. CRH^PVN^ neurons are activated in response to aversive stimuli^14–16^ which in turn drives CRH secretion from the median eminence, activation of pituitary corticotroph cells and the subsequent release of adrenocorticotropic hormone (ACTH). ACTH travels to the adrenal gland to stimulate de novo synthesis and secretion of corticosteroids. While ultradian rhythms in CRH secretion have been observed previously^17–21^, the reported period of these rhythms is variable ranging from minutes to hours. In addition, other studies show that ultradian pulses of corticosteroid secretion can be generated even if CRH secretion is clamped at a constant level^22^. Therefore, questions remain as to whether ultradian patterns of CRH^PVN^ neuron activity exist and how they correlate with ultradian corticosteroid secretion.

In addition to projections to the median eminence to control the HPA axis, CRH^PVN^ neurons also have axonal projections to other brain nuclei^23–27^. These central projections are important for the regulation of arousal and stress-associated behaviours^28^. While there is a large body of data showing that both CRH peptide or corticosteroids regulate arousal^29^, only recently has a direct role for CRH^PVN^ neurons been demonstrated. Specifically, optogenetic activation of CRH^PVN^ neurons was shown to promote wakefulness via projections to lateral hypothalamus orexin neurons^26,27^. Likewise, chemogenetic inhibition or ablation of CRH^PVN^ neurons decreased locomotor activity and wakefulness^26^. Using fiber photometry recordings of CRH^PVN^ neuron activity, one study observed increases in neural activity coincident with shifts from sleep to wake^26^. However, another investigation using similar methods did not observe this coincident activity^27^. One limitation of these studies is that they only analyzed CRH^PVN^ neural activity over time frames of seconds to minutes, precluding the detection of ultradian rhythms that manifest over longer periods. What is also unclear from previous work is how spontaneous CRH^PVN^ neuron activity is coordinated with both behaviour and ultradian pulses of corticosterone secretion.

## Results

### GCaMP6s fiber photometry recordings reveal ultradian rhythms in CRHPVN activity

To record the activity of the CRH^PVN^ neuron population in non-stressed freely behaving mice, we used GCaMP6s fiber photometry (Figure 1 a-c; see methods). On the day of recordings, mice were connected to the photometry system and then left undisturbed for 24 hours with free access to food and water in their home cage while CRH^PVN^ GCaMP6s activity was recorded along with behaviour with an overhead camera.

**Figure 1:**
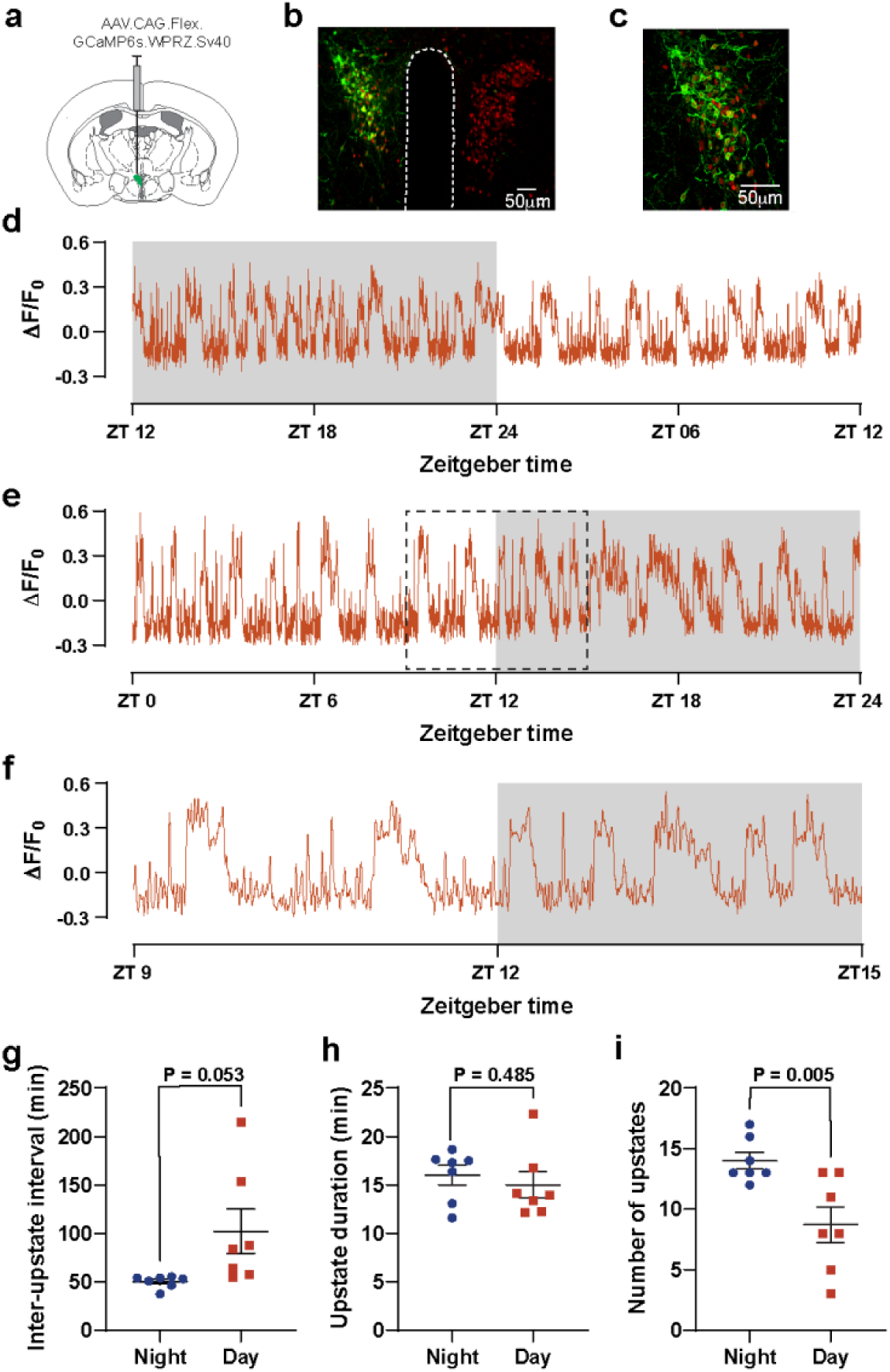
Ultradian rhythms of CRH^PVN^ neuron activity in freely behaving mice. a) GCaMP6s was targeted to CRH^PVN^ neurons via stereotaxic injection. b) Fluorescence image showing the expression of GCaMP6s (green) in CRH neurons (red) within the PVN. c) Higher magnification confocal image demonstrating co-localization of GCaMP6s (green) with CRH neurons (red) in the PVN. d/e) Example photometry recordings of CRH^PVN^ activity showing spontaneous ultradian pulses upstates 24-hours. Light off at ZT12, light on at ZT24. Dotted box in e is shown expanded in (f). g) Inter-upstate interval (min) shown for night versus day. Paired t test, t (6) = 2.404, p = 0.0530. h) Upstate duration (min) comparison between night and day. Paired t test, t (6) = 0.7443, p = 0.4848. i) Number of upstates observed compared between night and day. Paired t test, t (6) = 4.301, p = 0.0051.

To allow for long-duration recordings without photobleaching, we scheduled the acquisition of photometry recordings with a 3sec on/7sec off (30%) duty cycle. We averaged the data collected during the “on” period such that we effectively sampled CRH^PVN^ population activity once every 10 seconds (0.1Hz). This approach revealed slow ultradian events which we termed “upstates” (Figure 1d-f). Over the 24-hour day, there were on average 22.7 ± 1.9 upstate events (n=7) with a mean duration of 15.5 ± 1.0 min each (n=7, Figure 1h). The frequency of these upstate events is similar to the frequency of ultradian corticosteroid pulses previously reported in rats^30,31^. The number of up-states was higher during the dark phase of the day/night cycle (14.0 ± 0.7) compared to the light phase (8.7 ± 1.5, n = 7, P=0.005, Figure 1i). In between as well as during upstates, there was ongoing neural activity, consistent with the basal activity previously reported in the CRH neuron population^16,32,33^. However, the low sampling rate used for these experiments precluded the analysis of these fast events.

### Coordination of spontaneous CRH^PVN^ activity and behavioural arousal

Past work has suggested that CRH^PVN^ neurons are involved in regulating the sleep-wake cycle and behavioural arousal^26,27^. To determine if there was any correlation between spontaneous CRH^PVN^ neuron activity and behavioural arousal in our recordings, we analysed head movement in our 24-hour video recording with DeepLabCut^34^ (Figure 2b). Movements occurred over the entire 24-hour day and ranged from small head movements while the mouse remained within their nest, to locomotion where the mouse moved around the entire cage. When we plotted head distance moved per 10 sec bin, we observed periods of high levels of movement that reoccurred over the day (referred to as bouts). We analyzed the number of bouts of head movement over the day/night cycle and found that there were more bouts of head movement during the dark phase (10.0 ± 0.4) compared to during the light phase (5.4 ± 0.5, n= 7, P = 0.0007). We aligned the onset or offset of each bout of movement with photometry. We observed that CRH^PVN^ activity increased at approximately the same time as a movement bout was initiated. Likewise, CRH^PVN^ activity decreased at approximately the same time as a bout of movement was terminated, however, remained slightly elevated for longer (Figure 2c-e).

**Figure 2:**
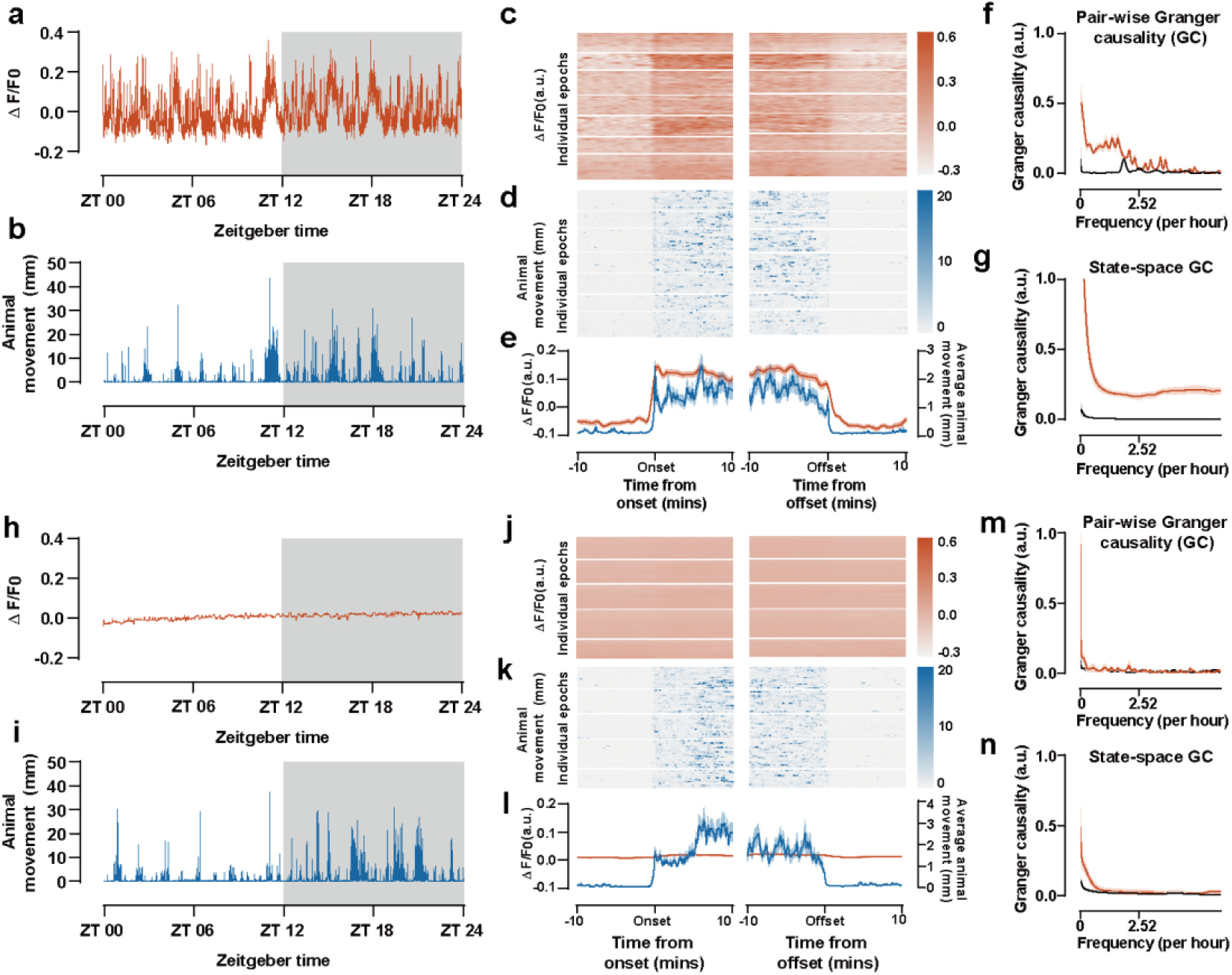
Correlation between CRH^PVN^ activity and behavioural arousal in mice. a) Example photometry recording showing spontaneous ultradian pulses across the 24-hour day. b) Corresponding 24-hour head movement profile (10sec bins). c) Heatmap of CRH^PVN^ activity before and after the onset and offset of movement bouts in 7 mice. d) Movement profiles corresponding to neuronal activity data shown in (c). e) Time-locked analysis of neuronal activity and movement around the onset and offset of a bout of movement. Red and blue lines represent average fluorescence and movement, respectively, aligned to the onset and offset of a bout of movement. f) Pair-wise Geweke Granger causality showing relative differences between estimated ΔF/F_0_ –> movement (red line) and movement –> ΔF/F_0_ (black line) causal influences in GCaMP6s expressing mice. g) State-space Granger causality showing relative differences between estimated ΔF/F_0_ –> movement (red line) and movement –> ΔF/F_0_ (black line) causal influences in GCaMP6s expressing mice. h) Example of photometry recording from a control mouse expressing GFP. i) Corresponding 24-hour movement profile for the mouse above. j) Heatmap of GFP fluorescence before and after the onset and offset of movement bout across 5 GFP-expressing mice. k) Movement profiles corresponding to data shown in above. l) Time-locked analysis of GFP ΔF/F_0_ and movement around the onset and offset of a bout of movement. Red and blue lines represent average fluorescence and movement, respectively, aligned to the onset and offset of a bout of movement. m) Pair-wise Geweke Granger causality showing relative differences between estimated ΔF/F_0_ –> movement (red line) and movement –> ΔF/F_0_ (black line) causal influences in GFP expressing mice. n) State-space Granger causality showing relative differences between estimated ΔF/F_0_ –> movement (red line) and movement –> ΔF/F_0_ (black line) causal influences in GFP expressing mice.

To investigate the temporal relationship between GCaMP6s CRH^PVN^ signals and movement, we performed pair-wise Geweke Granger and state-space Granger causality (GC) analysis (Figure 2f/g). These test whether one time series (i.e. GCaMP) is useful in forecasting another (i.e. movement). This analysis revealed that ΔF/F_0_ was more predictive of movement (orange line in Figure 2f and g) than in the opposite direction (movement predicting ΔF/F_0_, black line). This suggests the direction of coupling is driven by changes in CRH^PVN^ neuron activity. Both types of analysis show a predominant causal influence of ΔF/F0 to movement, particularly at lower frequencies. Similar results were observed using Convergent Cross Mapping analysis (data not shown). Together, these data show that changes in CRH^PVN^ neuron GCaMP6s fluorescence are predictive of animal activity, but not vice-versa.

As a control experiment, we injected a Cre-dependent GFP-virus into the PVN of Crh-Ires-cre mice and implanted an optic fiber probe as in the GCaMP6s mice. We then recorded photometry signals and behaviour using the same protocols as above. In these recordings, there was no evidence of distinct events or upstates (Figure 2h) despite bouts of animal movement (Figure 2i). We next repeated the pair-wise Geweke and state-space GC analysis (Figure 2m/n). For GFP recordings, since there is no fluctuation of fluorescence coupled to neural activity, the ΔF/F0 essentially reflects noise and background fluorescence. Consistent with this, the pair-wise Geweke and state-space GC estimates are low and do not indicate a preference for either direction. This suggests that the relationship observed in the GCaMP6s recordings is due to neural activity and not movement artifact.

### Chemogenetic activation of CRH^PVN^ neurons leads to elevations in home cage activity

We showed that CRH^PVN^ neuron activity precedes and predicts increases in behavioural arousal. We next tested if artificial activation of CRH^PVN^ neurons could drive increases in home cage behaviour. *Crh-Ires-Cre* mice were unilaterally injected with AAV9-hSyn-DIO-hM3Dq-mCherry to drive the expression of the excitatory DREADD receptor in CRH^PVN^ neurons (Figure 3a). After a minimum of 4 weeks, mice were habituated to handling and the presence of a rodent activity detector (RAD) device^35^ on their home cage. On the day of testing, mice were given either a single injection of vehicle or deschloroclozapine (DCZ) and home cage activity was recorded for two hours. Vehicle-injected mice showed a transient increase in home cage activity that returned to baseline by one hour post-injection (Figure 3b). DCZ injected mice, however, showed a much larger and sustained increase in home cage activity that persisted for the entire two hours of recording post-injection (Figure 3b). We compared the mean level of home cage activity between baseline (2 to 1 hours before injection) with 1 to 2 hours following injection (Figure 3c). While home cage activity was not different pre-versus post-injection for vehicle-treated mice, it was elevated for DCZ injected mice. A two-way ANOVA revealed a significant effect of DCZ treatment (F_(1,19)_=27.31, P <0.001), a significant effect of time (F_(1,19)_=35.08, P < 0.001) and a significant interaction (F_(1,19)_=26.54,P<0.001). This data suggests that activation of CRH^PVN^ neurons in the home cage environment can by itself drive increases in behavioural arousal in mice.

**Figure 3:**
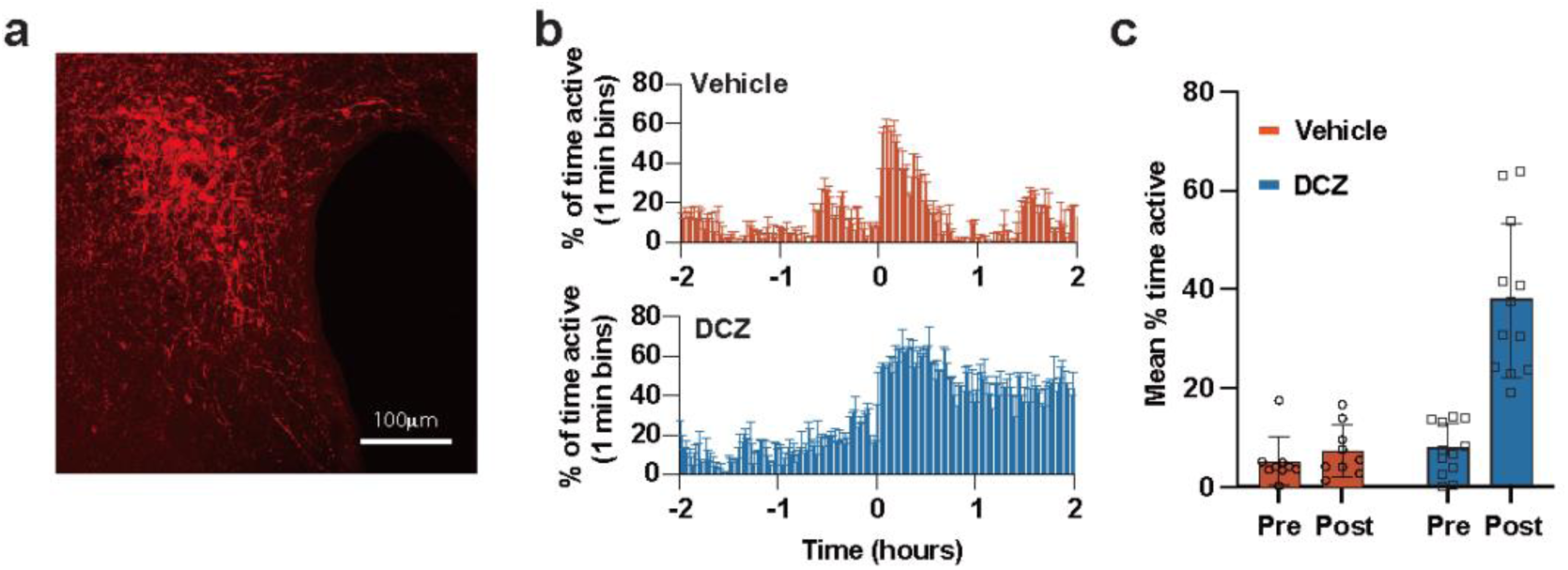
Chemogenetic activation of CRH^PVN^ neurons increases home cage activity. a) Image showing expression of hM3Dq-mcherry in the PVN. b) Mean ± SEM activity profiles (percentage of time active per 1-minute bin) for mice injected with vehicle (red) or DCZ (blue). Mice injected with vehicle (at time = 0) showed a transient increase in activity, whereas mice injected with DCZ showed a prolonged increase in activity. c) Summary data showing Mean ± SEM activity pre-injection (–2 to –1 hours) and post-injection (+1 to +2 hours).

### Coordinated CRH^PVN^ activity and behaviour in Crh-IRES-Cre rats

The above data show a clear relationship between spontaneous CRH^PVN^ activity and spontaneous behaviour in mice over the 24-hour day. The period of CRH^PVN^ activity over the 24-hour day appeared similar to the period of ultradian corticosterone pulses previously reported in rats^22,31^. To investigate this further, we generated a *Crh-Ires-Cre* rat so that we could perform CRH^PVN^ fiber photometry, behavioural analysis and repetitive blood sampling.

*Crh-Ires-Cre* rats were generated using CRISPR-Cas9 technology. Cre recombinase was expressed under the control of the endogenous *Crh* gene by inserting an internal ribosomal entry site (IRES)-Cre cassette immediately after the translation stop site of the *Crh* gene. We observed *Cre* mRNA labelling in brain regions where *Crh* is known to be expressed, including in the PVN (Supplementary Figure 1 and Supplementary Table 1). Immunohistochemistry for cre-recombinase protein showed labelling in the PVN of *Crh-Ires-Cre* rats (Supplementary Figure 2). To determine the exact colocalization of *Crh* and *Cre* mRNA, we performed dual-label RNAscope in the PVN. This showed that on average 81.9 ± 3.9 % of *Crh-*labelled neurons coexpressed *Cre* and that 97.8 ± 0.5 % of *Cre* labelled neurons coexpressed *Crh* (Figure 4a/b). To determine if knock-in of *Ires-Cre* affected basal corticosterone levels, we measured morning and evening corticosterone with tail vein blood sampling. We observed a clear elevation of corticosterone levels in the PM samples compared to AM samples of both wildtype and *Crh-Ires-Cre* rats. There was a significant effect of time of day (F_(1,3)_=23.55, P=0.0003), but no effect of genotype (F_(1,14)_=3.329, P=0.0895) and no significant interaction (F_(1,13)_=0.7100, P=0.4147; Supplementary Figure 2). Likewise, adrenal weight was not significantly different between wildtype and *Crh-Ires-Cre* rats (P=0.4701; data not shown). We next injected Cre-dependent GCaMP6s AAV into the PVN of *Crh-Ires-Cre* rats. To determine if GCaMP6s were targeted to CRH neurons, we performed RNAscope for *Crh* with immunohistochemistry for GCaMP. 58.34 ± 0.075% of *Crh* positive neurons expressed GCaMP6s while 87.71 ± 0.036% of GCaMP6s expressing neurons co-expressed *Crh* mRNA (Supplementary Figure 2).

**Figure 4:**
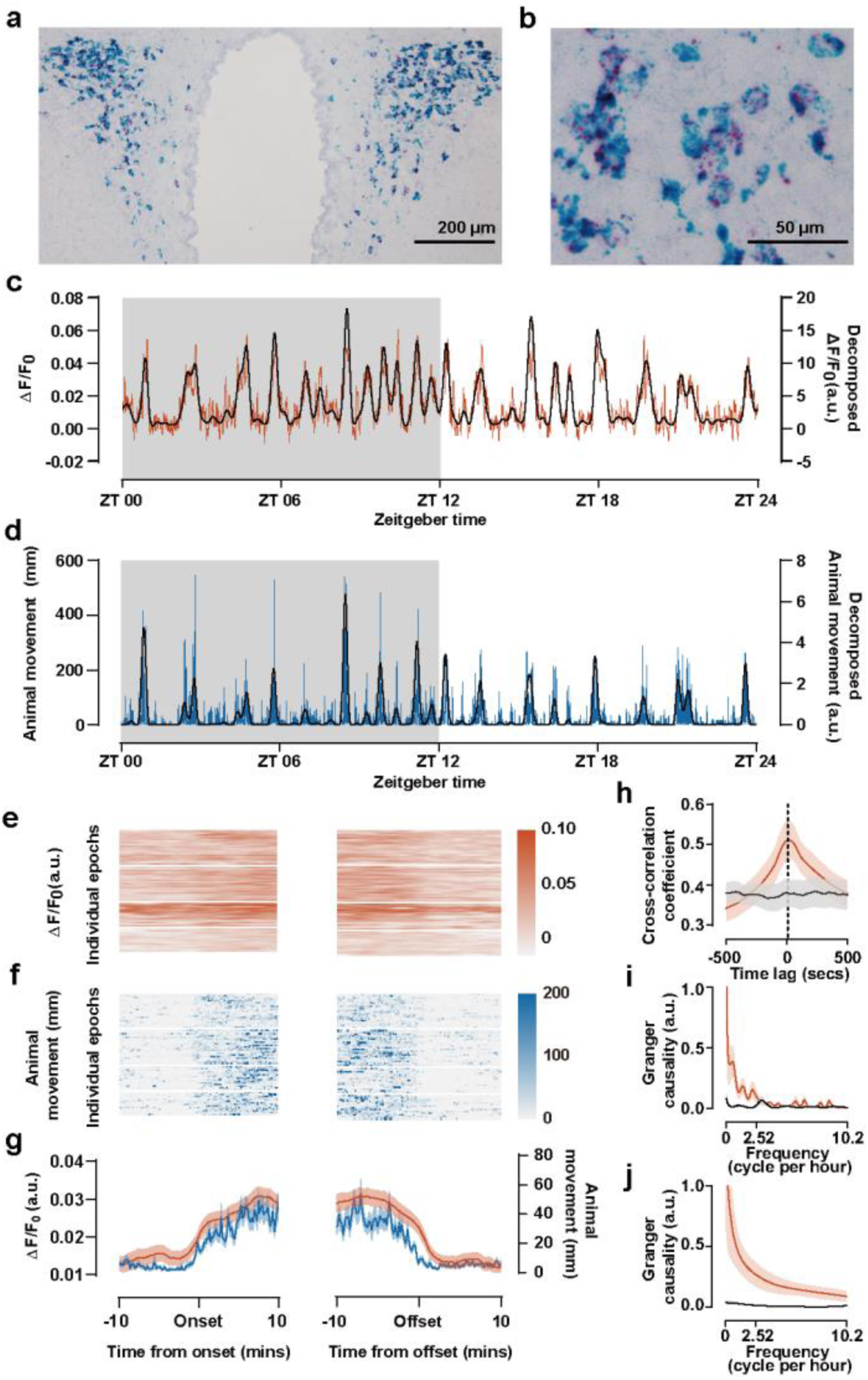
Spontaneous elevations in CRH^PVN^ activity correlate with bouts of behavioural arousal in *Crh-Ires-Cre* rats. a/b) Example images of RNAscope dual in situ hybridization in the PVN region for *Cre* (blue) and *Crh* (red) mRNA. c) Example photometry recording of CRH^PVN^ activity showing spontaneous ultradian activity across the 24-hour day in a freely behaving rat. Black line shows the slow component of the photometry recording extracted using decomposition. d) Simultaneous animal movement tracking from the same rat in (c). The black line shows the slow component of the animal movement tracking. e) Heatmap representation of CRH^PVN^ activity before and after the onset and offset of movement bouts from 4 GCaMP-expressing rats. f) Heat map movement profiles corresponding to neuronal activity data shown in (e). g) Average CRH^PVN^ activity (red) and animal movement (blue) 10 mins before and after the onset and offset of each movement bout. h) Cross-correlation analysis demonstrating a peak correlation coefficient of 0.51 (red line) at 10 secs (CRH^PVN^ activity preceding animal movement). No clear peak was observed in the randomly circular shifted data (black line) Granger-Geweke causality showing relative differences between estimated ΔF/F_0_ –> movement (red line) and movement –> ΔF/F_0_ (black line) causal influences in rats. j) the same as in (i) but with state-space Granger causality estimates of directional causal influence.

We next performed PVN fiber photometry on GCaMP6s expressing *Crh-Ires-Cre* rats over 28 hours along with video recording of behavior. As with recordings in mice, we ignored the first four hours after connecting the optic fiber. Similar to recordings from mice, we observed recurrent upstates of CRH^PVN^ neuron activity in rats that had a frequency of 1.0 ± 0.9 events per hour (n=4, Figure 4c). However, we observed that the magnitude of signals was smaller in rats compared to mice and upstates were less obvious. We next analysed head movement in our 24-hour video recording with DeepLabCut. Consistent with our observations in mice, bouts of head movements showed an ultradian rhythm that appeared coordinated with upstates of CRH^PVN^ activity. We aligned the onset or offset of a bout of movement with photometry and observed a similar relationship to that seen in mice (Figure 4d-g). Cross-correlation between ΔF/F_0_ and head movement revealed a peak correlation coefficient of 0.51 (red line) at 10 secs (CRH^PVN^ activity preceding animal movement). No clear peak was observed in control (randomly circular shifted) data.

To further investigate the temporal relationship between GCaMP6s CRH^PVN^ signals and movement in rats, we performed pair-wise Geweke and state-space GC analysis. Similar to what we observed in mice, the orange line (ΔF/F_0_ predicting movement) is consistently higher than the other direction (movement predicting ΔF/F_0_; black), confirming the direction of coupling is driven by changes in CRH^PVN^ neuron activity (Figure 4i/j).

To measure pulsatile corticosterone secretion, we performed jugular vein cannulation in *Crh-Ires-Cre* rats. After recovery from surgery, animals were connected to a custom-built automated blood sampling system. This system could automatically take 25-40 μL blood samples once every 7.5 or 15 min. Following collection of each blood sample, the same volume of saline was automatically infused back into the rat. Using this system, we were able to collect blood samples and detect spontaneous ultradian corticosterone pulses with temporal kinetics similar to that previously reported^30,31^ (Supplementary Figure 3). In a separate cohort, we performed jugular vein cannulation in animals that had previously undergone surgery for GCaMP6s fiber photometry. We then measured CRH^PVN^ activity, behavior and blood corticosterone over a 6-8 hr time window in freely behaving *Crh-Ires-Cre* rats. As can be seen from the representative recordings, bursts of CRH^PVN^ activity (Figure 5a/d) preceded a pulse of corticosterone (Figure 5 b/e) in many instances. However, in other instances, bursts of CRH^PVN^ activity occurred without any subsequent corticosterone pulse. There were also cases where corticosterone pulses occurred without clear bursts of CRH^PVN^ activity. We next performed cross-correlation to assess the relationship between CRH^PVN^ activity and corticosterone. On average, the cross-correlation coefficient between CRH neuronal activity and corticosterone secretion peaked at –15 min time lag with a coefficient of 0.73 ± 0.05 (n=4). This suggests that CRH neuronal activity and corticosterone secretion are correlated on average and that CRH^PVN^ activity precedes corticosterone by 15 min (Figure 5g). CRH^PVN^ activity and animal movement were also highly correlated, with a peak at 0 min time lag (Figure 5h).

**Figure 5:**
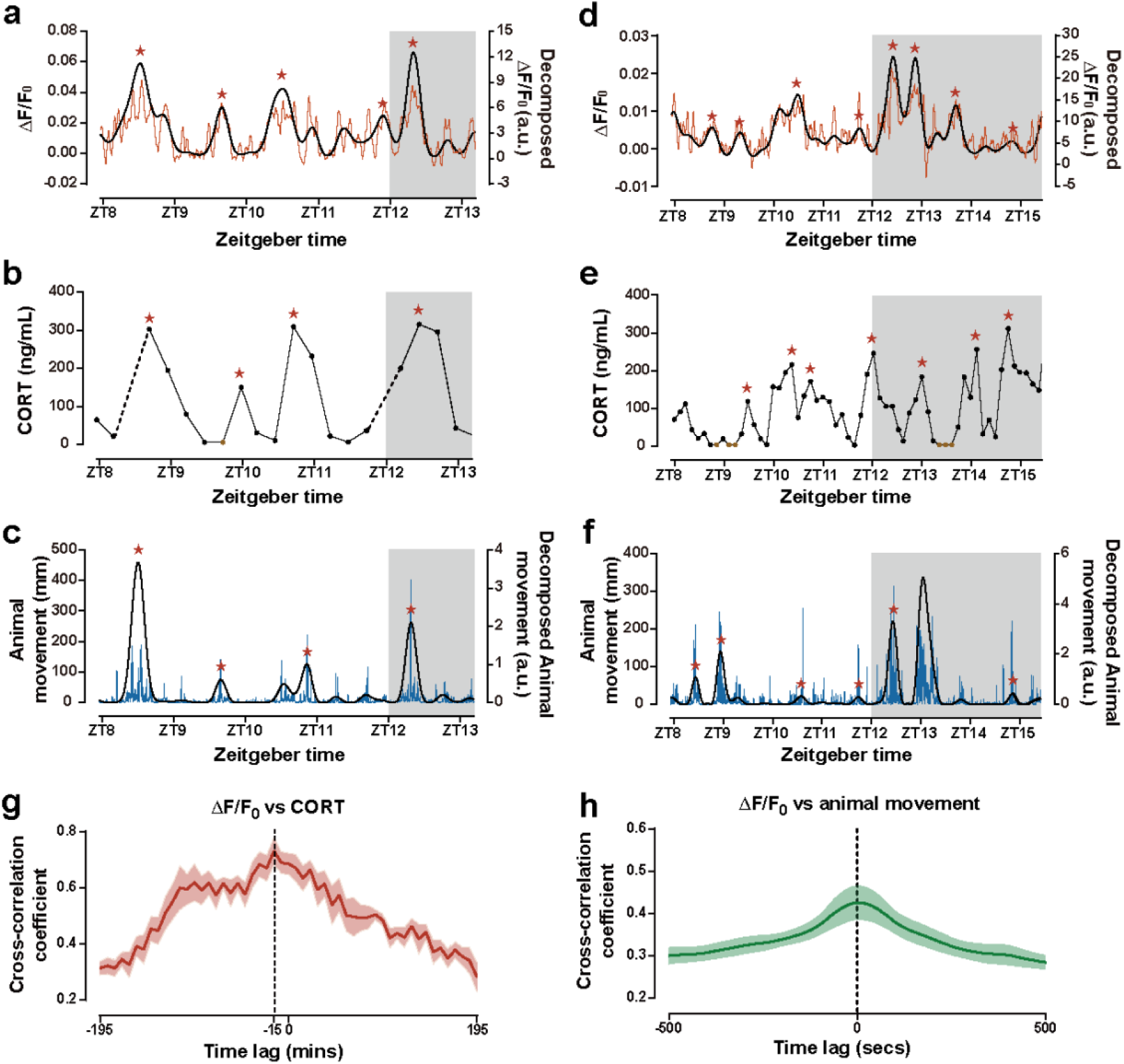
Ultradian CRH^PVN^ activity correlates with pulsatile corticosterone secretion under unstressed conditions. a) Example photometry recording of CRH^PVN^ activity with simultaneous blood sampling for corticosterone (CORT) (b) and recording of rat movement (c). Ultradian rhythms can be seen in all three measures. Black line shows the slow component of the photometry recording extracted using decomposition. Dashed line in b show missing samples (sampling rate of 1 sample/15 min). Red stars indicate ultradian pulses identified using MATLAB. d) Example photometry recording of CRH^PVN^ activity from a different rat along with simultaneous blood sampling for corticosterone (e) and recording of rat movement (f). Blood sampling in (e) was performed at a rate of 1 sample every 7.5 mins. g) Cross-correlation analysis of CRH^PVN^ neuron activity and CORT secretion show a peak correlation coefficient of 0.73 at –15 mins (CRH^PVN^ activity precedes corticosterone secretion). h) Cross-correlation analysis of simultaneous CRH^PVN^ activity and animal movement show a peak correlation coefficient of 0.43 at 0 mins.

## Discussion

Here we show that CRH^PVN^ neurons exhibit an ultradian rhythm in excitability over the 24-hour day. Increases in CRH^PVN^ activity were predictive of increases in animal movement and chemogenetic activation of these neurons increased locomotion/movement suggesting that these neurons are important in controlling arousal state. In a new line of *Crh-Ires-Cre* rats, we found that pulses of corticosterone secretion were sometimes, but not always, preceded by elevations of CRH^PVN^ neural activity. However, correlation analysis revealed that on average, increases in CRH^PVN^ activity preceded corticosterone rises by approximately 15 min. Together these data reveal that CRH^PVN^ neurons display an ultradian rhythm of activity over the 24-hour day that is coordinated with behavioural arousal, however, the relationship between CRH^PVN^ activity and pulses of corticosteroid secretion is not one-to-one.

A key finding from this study is the high correlation between CRH neuronal activity and movement under resting, unstressed states. Granger causality analyses indicated that CRH^PVN^ neuron activity predicted changes in physical activity. This suggests that CRH^PVN^ neurons might be an important node in the complex network that governs arousal state. This is in line with previous work showing that CRH^PVN^ neurons are involved in the regulation of arousal/wakefulness^26,27^. CRH^PVN^ neurons also synthesise glutamate and other studies that have manipulated PVN vglut neurons have observed similar results^36,37^. In addition, past work has also shown that CRH^PVN^ neurons regulate stress-related behaviours and affective states^14,15,23^. CRH^PVN^ neuron projections to the lateral hypothalamus appear to be particularly important for many of these actions^23,26,27^. However, CRH^PVN^ neurons have projections to other brain regions including the amygdala and periaqueductal gray^25^ and these projections are also likely to be important.

The most extensive CRH^PVN^ neuron projection is to the external zone of the median eminence where these neurons release CRH peptide to control the HPA axis^38^. Despite these neurons being widely accepted as master controllers of the HPA axis, there is still debate regarding their role in controlling the pulsatile, ultradian pattern of corticosteroid secretion. This debate has stemmed from past experiments which have shown that ultradian corticosteroid pulses can persist following disconnection of the hypothalamus from the pituitary in sheep^39^ or that corticosterone pulses can be generated with constant CRH infusions in rats^22^. This has led to the proposal of a sub-hypothalamic pulse generator mechanism for pulse generation^22,40^. Prior to this study, it was unknown whether the CRH^PVN^ neuron population exhibited ultradian rhythms of activity *in vivo* that are coordinated with corticosteroid release. Here, we demonstrate that CRH^PVN^ neurons do exhibit ultradian rhythms in activity over the 24-hour day in both rats and mice. In rats, we demonstrate that in many cases, these bursts of activity precede pulses of corticosterone release. Cross-correlation analysis also demonstrated a correlation between CRH^PVN^ activity and corticosterone with the peak in CRH^PVN^ neuron activity preceding corticosterone by 15 min. These data strongly suggest that bursts of CRH^PVN^ neuron activity contribute to ultradian pulses of corticosterone release. However, as noted above, the relationship is not one-to-one. This could be due to several reasons. Firstly, we recorded CRH^PVN^ neuron activity from only one side of the PVN. It is possible that some pulses of corticosterone release were driven by bursts of activity from the unmonitored contralateral PVN. Secondly, it is likely that ultradian patterns of CRH^PVN^ neuron activity cooperate with a sub-hypothalamic pulse generator mechanism which leads to the final pattern of corticosteroid secretion. Future work using both mathematical models^40^ and artificial manipulations of CRH^PVN^ neuron activity will be needed to address this.

Unlike circadian rhythms, ultradian rhythms do not appear to be controlled by a central master clock. Instead, the circuit mechanisms appear to be unique to each system^41,42^. Past work has shown that ultradian pulses of corticosterone persist when animals are exposed to constant light as well as following ablation of the suprachiasmatic nucleus^43^. While the circuit mechanisms controlling ultradian activity of CRH^PVN^ neurons and corticosterone are not clear^8^, the sub-paraventricular zone (SPZ) has been implicated as a possible region important for generating these rhythms^44^. Specifically, Wu *et al.* observed ultradian rhythms of activity in PVN and SPZ neurons using Ca^2+^ imaging in cultured brain slices^44^. These rhythms were blocked with tetrodotoxin and glutamate receptor antagonists suggesting that they require local network activity. However, the detailed circuit mechanisms driving these rhythms remain unresolved.

In summary, we have demonstrated that the CRH^PVN^ neuron population displays an ultradian rhythm of activity over the day-night cycle. Spontaneous elevations of CRH^PVN^ neuron activity are predictive of increases in behavioural arousal. We also show a temporal correlation between elevations of CRH^PVN^ activity and pulses of corticosterone secretion, with CRH^PVN^ activity preceding corticosterone. However, CRH^PVN^ neuron activity and pulses of corticosterone are not one-to-one. Together, these data revel the patterns of CRH^PVN^ neuron activity over the 24-hour day and their complex relationship with behaviour and stress hormone secretion.

## Materials and Methods

### Animals

All animals were housed under a 12h light/dark cycle in individually ventilated cages with ad libitum access to food and water. All Experiments were conducted in accordance with the New Zealand Animal Welfare Act and approved by the University of Otago Animal Welfare and Ethics Committee.

### Stereotaxic surgery for mice

Adult (10-12 week old) male *Crh-Ires-Cre*^45^ or *Crh-Ires-Cre;Ai14* (tdTomato reporter) mice^46^ were anesthetized with 2% isofluorane and placed in a stereotaxic frame. Adeno-associated virus (AAV) encoding GCaMP6s (AAV1.CAG.Flex.GCaMP6s.WPRE.SV40) or GFP (AAV9.Syn.DIO.EGFP.WPRE.hGH) was stereotaxically injected unilaterally into the PVN via a Hamilton syringe (–0.8mm AP, –0.25mm ML, –4.5mm DV) at a volume of 1μL over 10 min (Figure 1a-c). A fiberoptic cannula (400μm core, 0.48 N.A.; Doric Lenses) was then implanted at the same coordinates and secured using adhesive dental cement. A separate cohort of mice received stereotaxic surgery and injection of AAV9-hSyn-DIO-hM3Dq-mCherry. All mice were given carprofen (5mg/kg) and lidocaine (2%) during surgery and allowed to recover for 4 weeks before experimental recordings.

### Fiber photometry for mice

Photometry recordings of GCaMP6s fluorescence were acquired using Synapse software and a RZ5P processor from Tucker-Davis Technologies (TDT, Alachua Florida) and optical components purchased from Doric Lenses^16,42^. Excitation LEDs (465nm blue and 405nm violet) were sinusoidally modulated at 211 and 531 Hz, respectively. Excitation wavelengths were relayed through a filtered fluorescence minicube (spectral bandwidth: 460-490nm and 405nm) to a 400 μm 0.48 NA fiberoptic cable connected to the mouse. Light power for the 465nm wavelength at the fiber tip was 35-45 μW. A single emission (filtered at 500-550nm) was detected using a femtowatt photoreceiver (2151, Newport) with a lensed fiber cable adapter.

On the day of recordings, mice were connected to the photometry system and then left undisturbed for 28 hours with free access to food and water in their home cage while CRH^PVN^ GCaMP6s activity was recorded along with behaviour with an overhead camera. Connecting mice to the photometry system involved handling, which is a stressor. For this reason, we excluded the first 4 hours of recordings following connection and only analysed the last 24 hours of photometry data.For these recordings, photometry data was acquired in a scheduled recording mode, where both LEDs (405 nm and 465 nm) were switched on for 3 sec and off for 7 sec. This scheduled recording mode was chosen to reduce photobleaching GCaMP6s over extended recording periods.

All experiments were conducted in the animal’s home cage, which was placed in a custom-made apparatus (40 cm length, 40 cm width, 40 cm height) with white walls and transparent lid. Mice were habituated to the testing room and apparatus for 7 consecutive days prior to experimental manipulations. On the day of recording, mice had their optic fiber implant attached to an optic cable attached to a swivel. They were then left undisturbed for 28 hours with free access to food and water in their home cage.

### Activation of CRH^PVN^ neurons with DREADDs

To measure homecage activity following the activation of CRH^PVN^ neurons with DREADDs, we used the rodent activity detector (RAD). The RAD device uses a passive infrared sensor to detect motion and outputs data as percentage time spent active per minute^35^. Mice were habituated for 7 days in which they were handled to mimic injection. On day 3 of handling habituation, a dummy detector (i.e. case without passive infrared sensor) was added to the cage to habituate the mouse to the detector. On day 6 of handling habituation the dummy detector was replaced with the RAD to record baseline activity. On day 8 mice were injected with either 100μL vehicle or DCZ (1mg/kg in vehicle). The mouse and cage were then returned to the rack and activity was recorded for 2 hours.

### Generation of Crh-Ires-Cre rats

A Sprague-Dawley Crh-Ires-Cre knock-in rat line (SD-*Crh^tm^*^1^(Cre)*^Kji^*)(RRID:RGD_401976372) was generated using CRISPR technology by Cyagen Biosciences Inc. Cre recombinase was expressed under the control of the endogenous *Crh* gene by inserting an internal ribosomal entry site (IRES)-Cre cassette immediately after the translational stop site of the *CRH* open reading frame.

### Immunohistochemistry and RNAscope

Rats were terminally anaesthetised with sodium pentobarbital. Upon loss of the pedal withdrawal reflex, rats were transcardially perfused with 0.9% heparinised saline solution followed by 4% paraformaldehyde (PFA). Brains were post-fixed in 4% PFA for 24 hours before being saturated sequentially with 10% sucrose, 20% sucrose and 30% sucrose. After which, the brains were sectioned coronally on a cryostat at 30 μm thickness for Cre recombinase immunohistochemistry and 12 μm thickness for *Cre* and *Crh* mRNA RNAscope. Slices for immunohistochemistry were stored in tris-buffered saline (TBS) at 4 °C before being transferred to cryoprotectant whereas slices for RNAscope were collected on Superfrost plus slides and were stored at –80 °C.

Immunohistochemistry was used to label Cre recombinase in the PVN region in *Crh-IRES-Cre* rats. Brain slices were incubated in ethylenediaminetetraacetic acid (EDTA) at 90°C for 15 mins and then thoroughly rinsed in TBS before being quenched using hydrogen peroxide (3% hydrogen peroxide, 40% methanol, 57% TBS). Next, slides were placed in blocking buffer (0.4% Triton-X + 0.25% bovine serum albumin (BSA) + 2% normal goat serum (NGS) in TBS) for 90 min and proceeded with primary antibody incubation in rabbit anti-Cre recombinase (G. Schütz, Heidelberg^47^) diluted 1:5000 in blocking buffer for 1 hour at room temperature and then 48 hours at 4°C. Slices were then washed extensively in TBS before being incubated with biotinylated goat anti-rabbit IgG (H+L) secondary antibody (ThermoFisher Cat# B-2770, RRID: AB_2536431,1:500 in TBS + 0.4% Triton-X+0.25% BSA) followed by incubation in Vectastain Elite A/B solution (Vector laboratories, PK6100). Cre labelling was visualised by a chromogenic reaction using nickel sulfate-3, 3’-diaminobenzidine chromogen solution. Slices were dried and mounted in dibutylphthalate polystyrene xylene (DPX) mounting medium and imaged with a bright field microscope.

RNAscope 2.5 HD Duplex assay (Advanced cell diagnostics Inc., USA) was performed according to the manufacturer’s protocol to co-label *Crh* and *Cre* mRNA in the brain (*Crh* probe Cat# 318931; *Cre* probe Cat# 423321). Brain slices were baked at 60°C for 30 mins and then incubated in pre-chilled 4% PFA for 15 mins before being dehydrated sequentially in 50%, 70% and 100% ethanol. Slides were incubated in RNAscope hydrogen peroxide for 10 mins at room temperature and then submerged in target retrieval solution at 98-102°C for 5 mins. Slides were then dehydrated in 100% ethanol and air-dried. Immedge™ hydrophobic barrier pen was used to draw a hydrophobic barrier around each slice. One drop of Protease Plus was added to each slice and then incubated for 30 mins at 40°C. Slides were washed in distilled water before being incubated with the appropriate probe for 2 hours at 40°C. After incubation, slides were kept in 5X saline-sodium citrate buffer overnight at room temperature.

On the following day, slides went through 6 hybridisation steps using hybridise Amp 1-6 solutions. After the sixth hybridisation step, Red solution mix containing 1 part of Red-B in 60 parts of Red-A was applied to each slice to detect the red signal (*Cre* mRNA). Four more hybridisation steps were performed before adding Green solution mix containing 1 part of Green-B in 60 parts of Green-A to each slice to detect green signal (*Crh* mRNA). Slices were thoroughly washed in Wash buffer and then washed in distilled water before being submerged in a 10% haematoxylin staining solution for 2 seconds. Slides then went through multiple wash steps in tap water, 0.02% ammonia water and then tap water again. Slides were dried at 60°C and then mounted using VectaMount mounting medium. The mounting medium was allowed to dry before slides were examined under a standard bright field microscope at 20-40x magnification. In the PVN, cells with only *Crh* mRNA expression, cells with only *Cre* mRNA expression and cells with both *Crh* and *Cre* mRNA expression were counted to calculate the co-expression levels. Cells were classed as expressing mRNA if they had three or more stained puncta.

To label for *Crh* mRNA and GCaMP6s protein, Crh-Ires-Cre rats that were previously injected with GCaMP6s-AAV were anaesthetised and perfusion fixed. Brain sections of the PVN were first processed with RNAscope Multiplex Fluorescent Detection kit v2 (Cat #323110) according to the manufacturer’s protocol. Day-1 slide processing was same as described above for RNAscope 2.5 HD Duplex assay. On day-2, three hybridization steps were performed at 40 °C in the hybridization oven using FL v2 Amp 1-3 solutions followed by three HRP steps including the addition of FL v2 HRP-C1 (Crh probe in C1 channel), fluorophore Cyanine 3 (Akoya BioSciences, # SKU NEL7444001KT) and FL v2 HRP blocker. Immunolabeling with GCaMP6s was continued immediately by pre-treating the sections with 4% cold PFA for 5 min and placing the slides in blocking buffer for 60 min followed by primary antibody incubation (chicken anti-GFP (1:2000) RRID: AB_10000240, diluted in blocking buffer) for 24 hours at 4 °C. The brain slices were thoroughly washed in TBS and then transferred into secondary antibody which contained goat-anti-chicken conjugated with Alexa Fluro-488 (RRID: AB_2543096) diluted 1:500 in TBS+ 0.4% Triton-X+0.25% BSA. The brain slices were then thoroughly washed in TBS before being cover slipped with Prolong Antifade Gold (ThermoFisher, P36930).

### Stereotaxic surgery for rats

Female heterozygous *Crh-Ires-Cre* rats (2-3 months old) were anaesthetised with isoflurane (5% in an induction chamber, and then 1-3% during the surgery) and placed in a stereotaxic frame. A 1 μl Hamilton syringe at –1.5 mm AP, +1.5 mm ML, –7.5 mm DV with a 10-degree angle was inserted into the brain to inject 0.7 μl of AAV1.CAG.Flex.GCaMP6s.WPRE.SV40 (PennVector Core, USA) at a 100 nl/min rate into the PVN. A fiber implant (400/430-0.48_8.5mm_MF1.25_FLT, Doric Lenses, Quebec, Canada) was lowered into the brain at a 10-degree angle (–1.6 mm AP, +1.7 mm ML, –6.8 mm DV) and secured using dental cement.

### Fiber photometry in rats

Optical recordings of GCaMP6s fluorescence were acquired using a custom software acquisition system with optical components purchased from Doric Lenses^16,42^. Excitation light-emitting diodes (LED) (465 nm and 405 nm) were sinusoidally modulated at 211 Hz and 531 Hz, respectively. Excitation wavelengths passed through a filtered fluorescence mini cube (spectral bandwidth: 460-490 nm and 405 nm) to the rat via a 400 μm, 0.48 N.A. optic patch cord. The light power of the 465 nm LED at the fiber tip was 35 μW, and the 405 nm LED was set to 15 μW light output. Emitted fluorescence was filtered (500-550 nm) and was collected onto a photoreceiver (2151, Newport).

Fiber photometry recordings were conducted in a custom round home cage (35cm in diameter, 40cm in height). All recordings started 4 hours before ZT12, and the rats were then left undisturbed for 28 hours. During the 28-hour recording, both 405 nm and 465 nm LED were set to turn on for 3 seconds and then to turn off for 7 sec to minimise bleaching.

### Photometry analysis

The isobestic 405 nm signal was scaled and fitted to the 465 nm signal using a second-order polynomial fit. The ΔF/F_0_ was calculated by subtracting the fitted 405 nm signal from the smoothed 465 nm signal and then dividing it the line of polynomial fit.

### Jugular surgery and automated blood sampling in rats

Female *Crh-Ires-Cre* (+/-) rats aged 3-4 months were used. Rats were anaesthetised with isoflurane (5% in an induction chamber, and then 1-3% during the surgery) and positioned in the dorsal position. A thermoplastic polyurethane elastomer catheter was inserted into the jugular vein through a skin incision. The jugular vein catheter was then tunnelled subcutaneously and exteriorised through an incision between the scapulae. The jugular vein catheter was filled with sterile heparinised saline (50 IU/mL) which was flushed twice a day.

Two different blood sampling protocols were used. For the first protocol, 1 sample was collected every 15 mins for 6 hours (24 samples). Each sample contained approximately 20 μL of saline and 40 μL of blood. The second protocol collected 64 samples across 8 hours (1 sample every 7.5 mins). Each sample contained approximately 20 μL of saline and 25 μL of blood. As the blood samples collected showed slight variation in dilution, different amounts of 0.9% saline (50 IU heparin per mL) were added to each sample to reach a common 1:10 dilution (1 part blood, 9 parts saline). The amount of saline required was determined by measuring the haematocrit of each sample.

### ELISA

Corticosterone concentrations were measured using the DetectX Corticosterone Enzyme Immunoassay kit (Arbor Assays, Ann Arbor, Michigan, USA Cat# K014, RRID AB_2877626) according to the manufacturer’s instructions. The assay sensitivity was 26.54 ng/mL and the intra-assay coefficient of variation for this set of experiments was 4.81%.

### Animal movement tracking

DeepLabCut was used to track animal movement^34,48^. The right and left ears of the mouse, the optic fiber cannula and the base of the tail were manually labelled in 25 frames chosen through k-means clustering. A ResNet-101 based neural network with default parameters was created and then trained until the loss plateaus for each different video resolution. The network was then used to analyse videos with specific resolutions, and the XY position of each body part was exported as a CSV file. In Matlab, we first normalised all coordinate data to the dimension of the highest input video resolution (1024 x 576 pixels) and took the median value between the ears and the cannula as a single tracking point. Large tracking jitters (> 20 px displacement between time points) were removed and interpolated XY coordinates were submitted to a 11 point median filter for smoothing.

Rat movements were also tracked using DeepLabCut. A ResNet-50 based neural network with default parameters was trained for 250,000 iterations on 19 manually labelled frames (left ear, right ear and fiber optic cannula were labelled) for each rat individually. The network was then used to analyse videos from that specific rat. The centroid of the labelled body parts was calculated. The displacement of the centroid was reported as animal movement in millimetres.

### Movement-triggered analysis

Since the tracking and photometry data are not sampled at the same rate, we downsampled tracking data to 0.1 Hz by removing the first second of recording and then averaging the 2 secs of recordings corresponding to the 3 secs of the photometry data. Using this downsampled tracking, multiresolution decomposition was done with a symlets 4 wavelet (sym4) at 6 levels. We summed the two fastest components to extract the sharp changes in the tracking data. To map these sharp transient points to bouts of activity, we also used the residual (slowest component) of the decomposition to exclude sustained activity bouts so that only fast, isolated bouts of activity are included in further analyses. The timestamps of these transitions were collected and used for movement-triggered analyses.

### Granger causality analyses

*Pair-wise Geweke Granger causality (GGC):* is based on Granger’s original idea that if a backwards time-shifted variable can better account for the variation in another, then the shifted variable “causes” the other in a statistical context (i.e., variance accounted for). The method for pairwise Granger’s causality was adopted from Cui et al., 2008^49^. In the current implementation, stepwise least squares auto-regressive models of time-synchronised tracking and photometry data are generated to obtain an optimal model order using the Schwarz’s Bayesian criterion (SBC; up to an order of 200). The frequency spectra modelling was generated by using a modified Levinson-Wiggins-Robinson (LWR) recursive algorithm, and the “causal influence” from one variable to another at specific frequencies was calculated as the mutual information between the variable itself normalised against the contribution from the other variable. A higher value between the two variables indicates a relatively minor contribution from the shared covariance, thus likely to “Granger cause” the other.

*State-space causality:* Downsampling by taking the median of windowed data results in the loss of information and likely exacerbates the moving average feature of our tracking and photometry data. To account for these potential biases, we also performed Granger causality (GC) analysis using a state-space model. The method for state-space Granger’s causality was adopted from Barnett and Seth, 2015^50^. Consistent with various generalizations of GC, linear state-space model uses optimal model order from SBC to estimate Kalman states from a singular decomposition weighted by future/past canonical correlation weights of the inputs (i.e., movement and photometry data). With defined (Kalman) states, the observation matrix, state transition matrix, innovations covariance matrix and Kalman gain matrix are derived to generate a state-space model in the innovations form. These parameters are used to generate error covariance matrices to parse out causal influences between frequency spectra calculated from an AR representation of the state-space model for tracking and photometry data. Causal influences from movement to photometry data, and vice versa, are determined by the relative mutual information after accounting for covariances from the other (as in GGC).

## Statistics

Statistical analyses were performed with Prism and MATLAB. Paired t-tests were used to analyse circadian difference in ultradian CRH neuronal activity. Two-way ANOVA was used to analyse the effect of DCZ treatment. All data are presented as mean ± SEM. *P < 0.05, **p < 0.01, and ***p < 0.001.

## Acknowledgments

This work was supported by a University of Otago Research Grant. SZ and CMBF were supported by University of Otago PhD scholarships.

## Data, Materials and Software Availability

All study data are included in the article and/or SI appendix.

## Figures and Tables

**Supplementary Figure 1:**
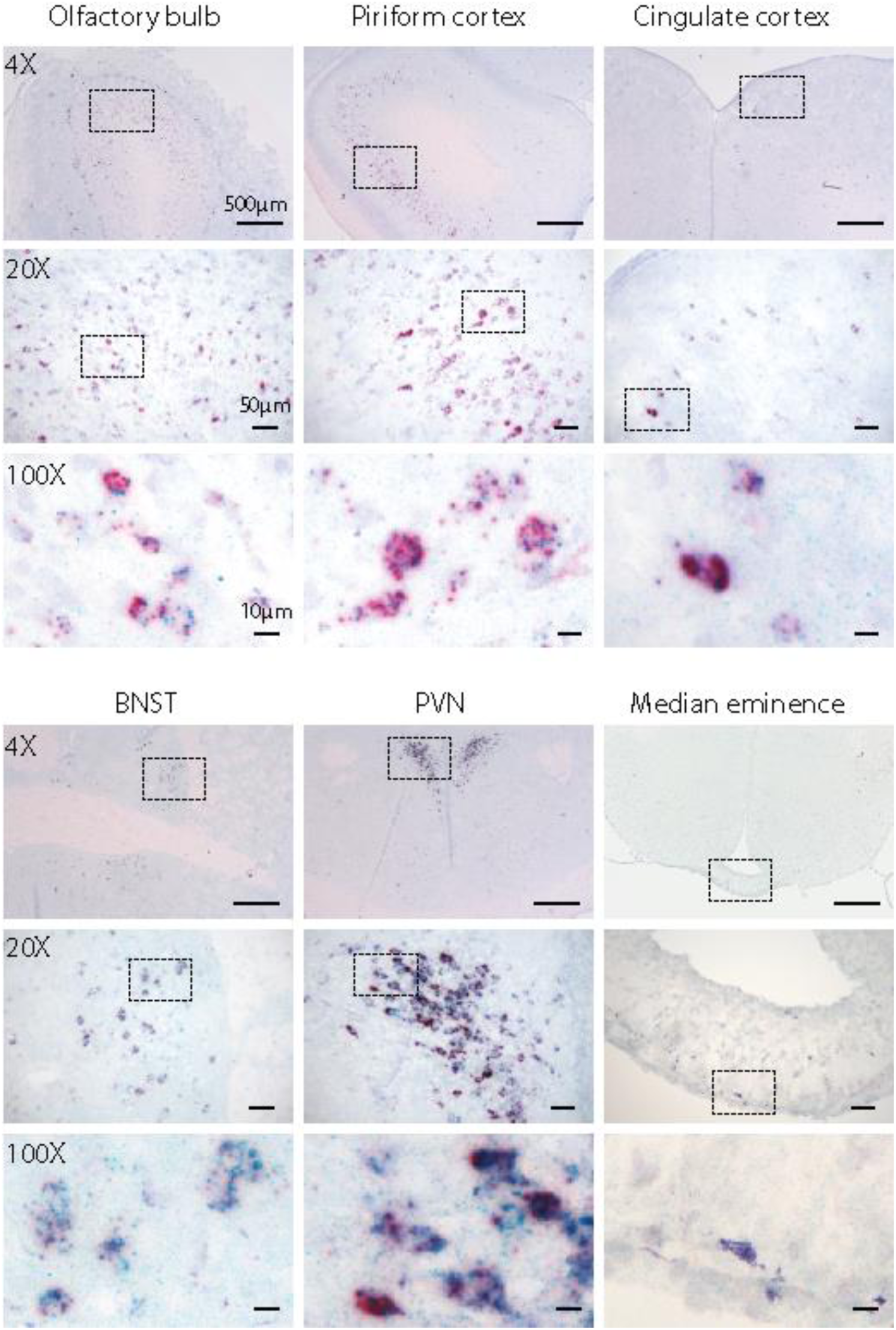
Co-localisation of *Crh* and *Cre* in *Crh-Ires-Cre* rats. RNAscope images displaying the co-localisation pattern of *Crh* mRNA (red) and *Cre* mRNA (blue) in various brain regions (olfactory bulb, piriform cortex, cingulate cortex, bed nucleus of the stria terminalis (BNST), PVN, and median eminence) in *Crh-Ires-Cre* rats. For each region, a progression of magnifications is shown (4X, 20X, and 100X).

**Supplementary Figure 2:**
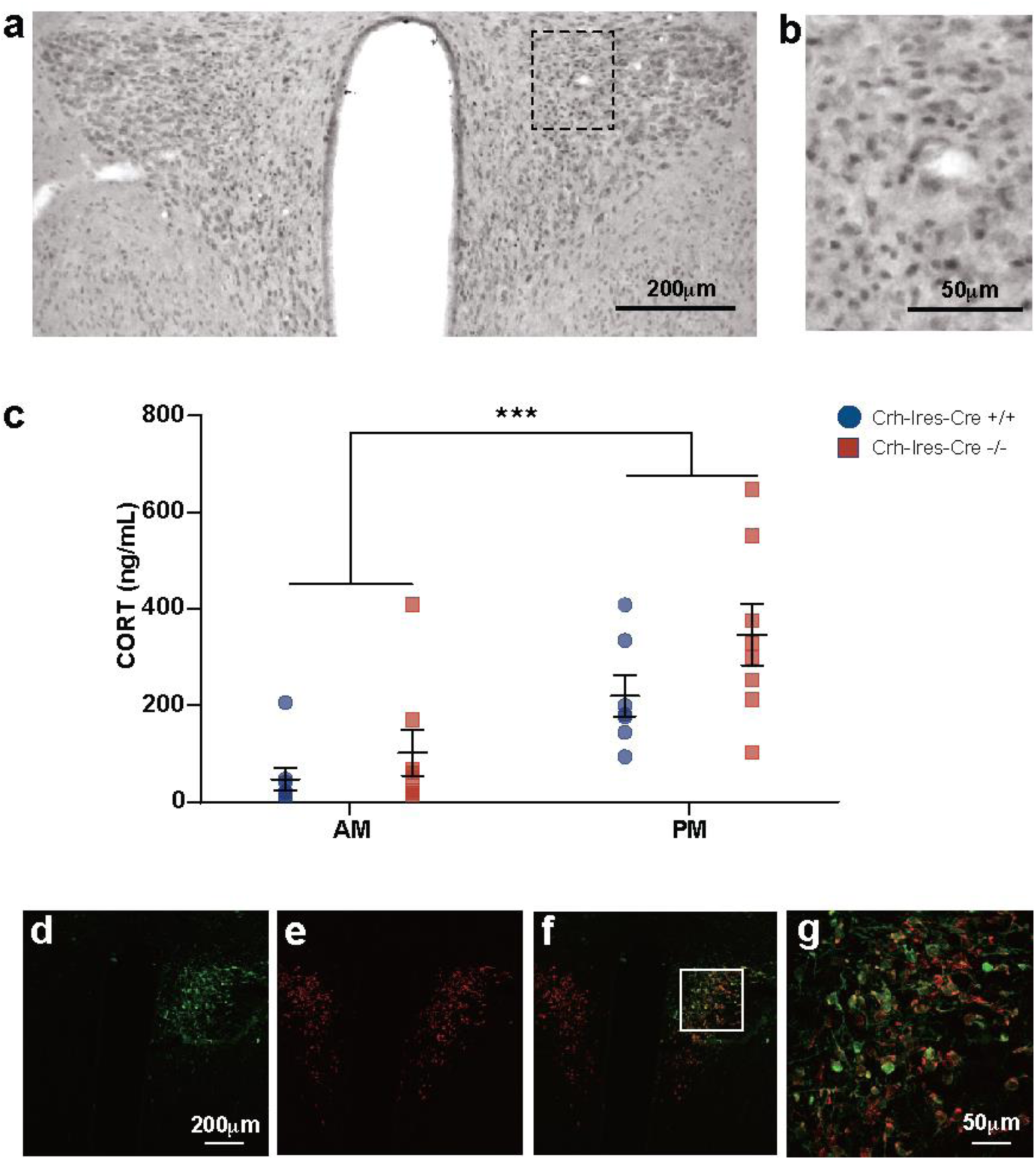
Characterization of Cre Recombinase expression and corticosterone levels in *Crh-Ires-Cre* rats. a) Example image of immunohistochemistry in the PVN region for Cre recombinase. Scale bar = 200 µm. b) Zoomed in section outlined in a. Scale bar = 50 µm. c) Mean ± SEM serum corticosterone (CORT) concentration (ng/mL) at AM and PM in *Crh-Ires-Cre* rats (in blue) vs wild type rats (in red). A mixed effect analysis revealed there was a significant effect of time of day (F (1, 3) = 23.55, p = 0.0003), but no significant effect of genotype (F (1, 14) = 3.329, p = 0.0895) and no significant interaction (F (1, 13) = 0.7100, p = 0.4147). d) Stereotaxic injection of GCaMP6s-AAV induces GCaMP6s expression in CRH^PVN^ neurons. *Crh* mRNA (red) is labelled on both sides of the PVN. e) GCaMP6s protein is labelled unilaterally on one side of the PVN. f) Merge of d and e showing co-expression of *Crh* mRNA and GCaMP6s. g) Zoomed-in images of the section outlined in f.

**Supplementary Figure 3:**
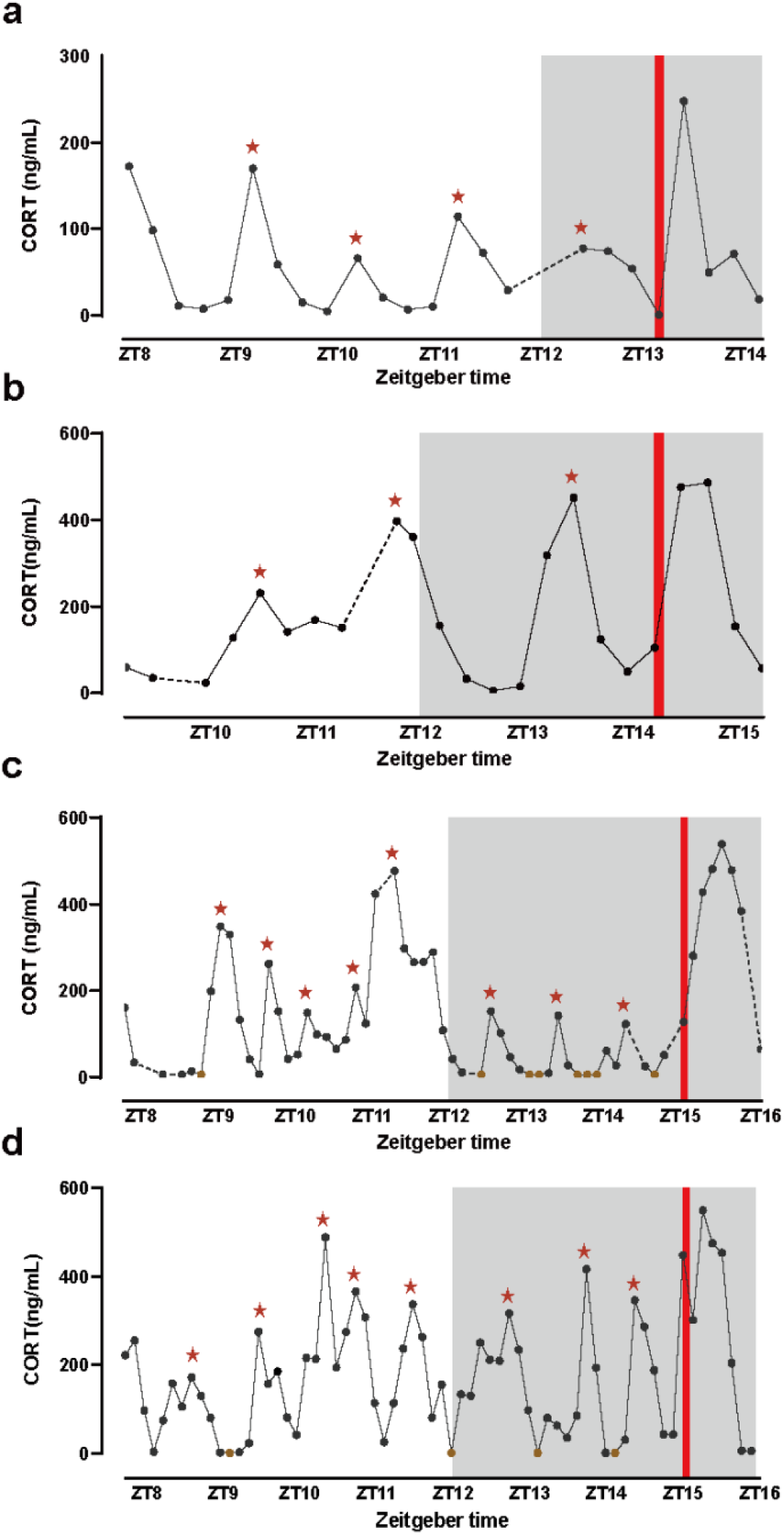
Corticosterone secretion under unstressed and stressed states. Jugular blood sampling in freely behaving rats reveals an ultradian rhythm in corticosterone (CORT) secretion. a) and b) shows ultradian peaks identified in CORT levels in 2 rats across 6 hours sampled once every 15 mins. c) and d) shows ultradian peaks identified in CORT levels in 2 rats across 8 hours sampled once every 7.5 mins. The red stars indicate ultradian peaks identified by Matlab. The red line indicates exposure to 5 mins of white noise stress (90dB). Dotted lines indicate missing value. Brown dots indicate samples with CORT values lower than the detectable range.

**Supplementary Table 1.**
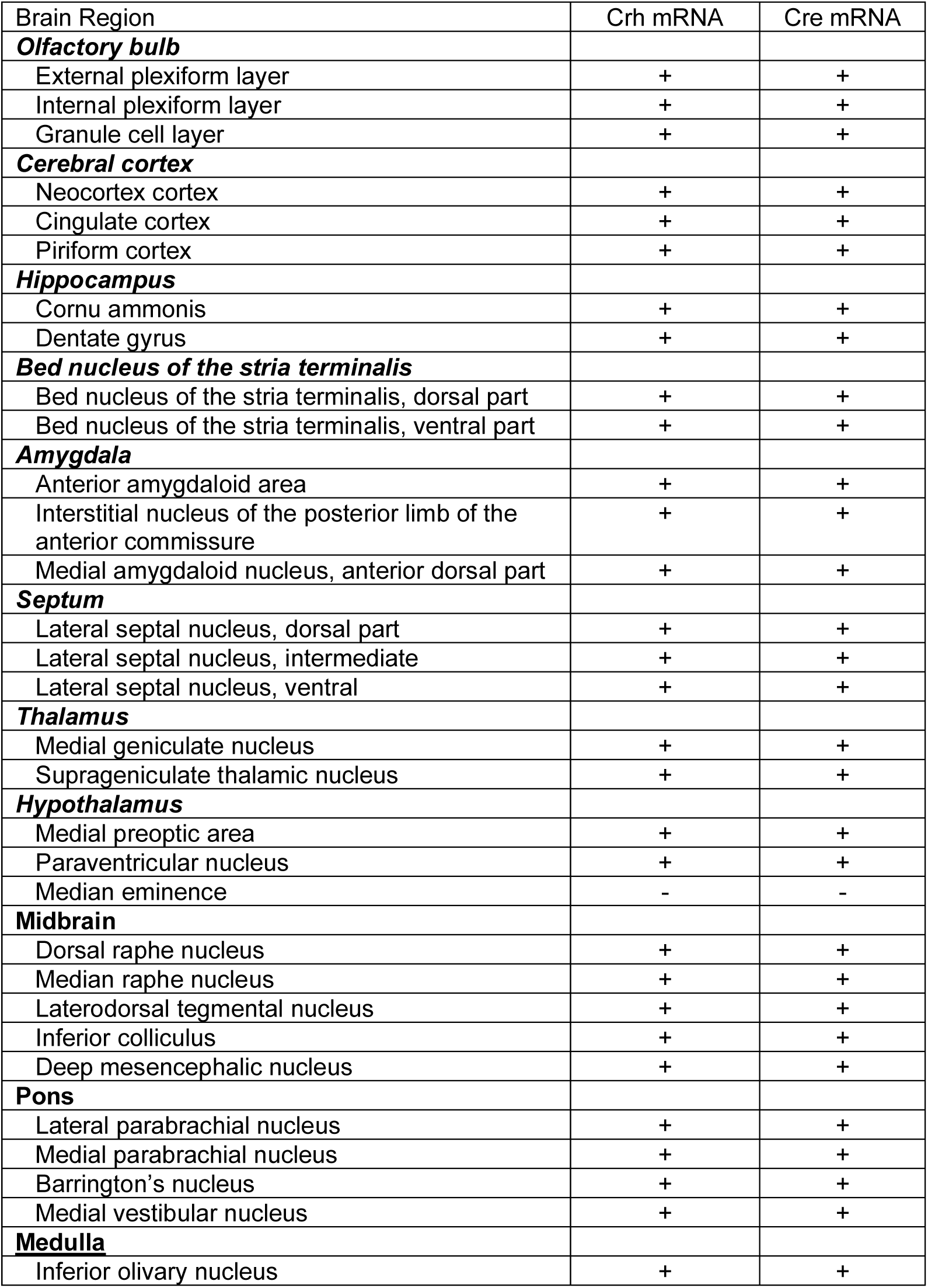
Co-expression of *Crh* and *Cre* mRNA across the brain in Crh-Ires-Cre rat brain tissue labelled with RNAscope. Example images shown in Supplementary Figure 1.

